# Involvement of Lateral Habenula Dysfunction in Repetitive Mild Traumatic Brain Injury-Induced Motivational Deficits

**DOI:** 10.1101/2022.05.04.490685

**Authors:** William J. Flerlage, Ludovic D. Langlois, Milan Rusnak, Sarah C. Simmons, Shawn Gouty, Regina C. Armstrong, Brian M. Cox, Aviva J. Symes, Mumeko C. Tsuda, Fereshteh S. Nugent

## Abstract

Affective disorders including depression (characterized by reduced motivation, social withdrawal and anhedonia), anxiety and irritability are frequently reported as long-term consequences of mild traumatic brain injury (mTBI)^1^ in addition to cognitive deficits, suggesting a possible dysregulation within mood/motivational neural circuits. One of the important brain regions that control motivation and mood is the lateral habenula (LHb) whose hyperactivity is associated with depression^2^. Here we used a repetitive closed head injury mTBI model that is associated with social deficits in adult male mice^3^ and explored the possible long-term alterations in LHb activity and motivated behavior 10-14 days post-injury. We found that mTBI increased the proportion of spontaneous tonically active LHb neurons while decreased LHb bursting. Additionally, mTBI diminished spontaneous glutamatergic and GABAergic synaptic activity onto LHb neurons, while synaptic excitation and inhibition (E/I) balance was shifted toward excitation through a greater suppression of GABAergic transmission. Behaviorally, mTBI increased the latency in grooming behavior in sucrose splash test suggesting reduced self-care motivated behavior following mTBI. To show whether limiting LHb hyperactivity could restore motivational deficits in grooming behavior, we then tested the effects of Gi (hM4Di)-DREADD-mediated inhibition of LHb activity in sucrose splash test. We found that chemogenetic inhibition of LHb glutamatergic neurons was sufficient to reverse mTBI-induced delays in grooming behavior. Overall, our study provides the first evidence for persistent LHb neuronal dysfunction due to an altered synaptic integration as causal neural correlates of dysregulated motivational states by mTBI.

## Introduction

Worldwide, millions of individuals are living with the long-term consequences of traumatic brain injury, making it one of the most societally and economically important issues today. The vast majority of traumatic brain injury cases are classified as mild. Mild traumatic brain injury (mTBI) occurs when mechanical energy is applied to the head with sufficient force to result in a temporary and spontaneously recovering alteration of consciousness lasting less than 24hrs, without the presence of brain imaging- indicated structural abnormalities ^4^. Incidence rates of mTBI within the United States have risen over the last decade, with estimates ranging from 500-800 per 100,000 population annually, a likely underestimate due to under-reporting. While the majority of mTBI cases post-concussive symptoms resolve within the days and weeks following injury, 15-25% of patients will continue to report physical, cognitive, and psychosocial disturbances in the months to years that follow ^1, 5^. There is also a growing appreciation of the compounded risk and severity of chronic psychiatric morbidity among those sustaining multiple head injuries, with the most at-risk populations include contact-sports athletes, military personnel, and victims of domestic abuse ^6, 7^. Chronic mood-related psychiatric morbidity following mTBI often presents as affective deficits (e.g., major depressive disorder, apathy) and associated dysregulation of emotional and social behaviors (irritability, aggression, suicidality, social withdrawal, anxiety) ^1^. Compelling evidence from animal and human studies suggest that long-term negative effects on social functioning and mood states are related to dysfunction of brain reward and motivational circuits including the lateral habenula (LHb)^2, 8, 9^. The LHb is a diencephalic structure that has emerged as an anti-reward hub for motivation and decision-making which links forebrain limbic structures with midbrain monoaminergic centers ^10-13^. LHb neurons receive glutamatergic, GABAergic and co-releasing glutamate/GABA inputs from the basal ganglia and diverse limbic areas including medial prefrontal cortex (mPFC), the internal segment of the globus pallidus (analogous to the rodent entopeduncular nucleus, EP), lateral preoptic area (LPO), lateral hypothalamus (LH), ventral pallidum (VP), medial and lateral septum, central amygdala (CeA), bed nucleus of stria terminals (BNST) as well as receiving reciprocal input from the ventral tegmental area (VTA). LHb projects to the substantia nigra, VTA, rostromedial tegmental area (RMTg), dorsal raphe nucleus (DRN), locus coeruleus (LC) and periaqueductal gray (PAG)^9, 14^. Promotion of LHb hyperactivity with excitatory synaptic inputs or dampening of inhibitory inputs mediates behavioral aversion, lack of motivation, anxiety and depressive phenotypes. Conversely, inhibition of LHb activity through inhibitory inputs or diminished excitatory drive reduces behavioral anxiety and depression, and promotes positive reinforcement ^9, 12, 15-20^. Specifically, LHb hyperactivity/ dysfunction is a common finding associated with anhedonia and social withdrawal which reflect some of the core features of reward and motivational deficits seen in depressed humans ^15, 21-23^. Not surprisingly, LHb represents a critical brain region that can be targeted for treatment of reward dysregulation in mood disorders ^19, 24-27^. However, whether LHb dysfunction plays a role in mTBI-induced psychopathology was unexplored. Therefore, we used a repetitive closed head injury mouse model of mTBI that has been shown to induce deficits in social interaction in adult male mice at 21 days post-injury ^3^ to test the effects of this mTBI model on LHb physiology and explore the possible involvement of LHb dysfunction in mTBI-induced changes in motivated self-care behavior in adult male mice at 10-14 days post-injury. This model of mTBI does not cause neuropathology; i.e., no neuroimaging evidence of micro-hemorrhages or overt damage to skull or brain while the mice show a brief loss of consciousness (delayed righting reflex). The model induces low levels of axonal damage in the corpus callosum and the cortex as well as astrogliosis and microglial activation, both expected to be observed in mTBI ^3^. Given that the medial cortical region under the impact site used in this mTBI model was likely to alter the function of subcortical regions including diencephalic structures such as the LHb that receive direct and indirect projections from cortical regions, we tested the effects of mTBI on LHb neuronal and synaptic function and self-care grooming behaviors. Here, we provide the first evidence for a causal link between mTBI-related LHb hyperactivity due to synaptic excitation/inhibition (E/I) imbalance in LHb neurons and motivational deficits in self-care and grooming behavior in adult male mice long after the initial injury. Our study presents possible therapeutic avenues for future interventional research directed at the LHb and related circuits for reversal of prolonged negative consequences of mTBI on mood, motivation and emotional regulation.

## Materials and Methods

### Animals

All experiments were carried out in accordance with the National Institutes of Health (NIH) *Guide for the Care and Use of Laboratory Animals* and were approved by the Uniformed Services University Institutional Animal Care and Use Committee. C57BL/6 mice (Charles River) were acquired at ∼postnatal day 35-49 (PN35-P49) and allowed at least 72hrs of acclimation before the initiation of any experimental procedures. Mice were group housed in standard cages under a 12hr/12hr light-dark cycle with standard laboratory lighting conditions (lights on, 0600-1800, ∼200lux), with ad libitum access to food and water. All procedures were conducted beginning 2–4hr after the start of the light-cycle, unless otherwise noted. All efforts were made to minimize animal suffering and reduce the number of animals used throughout this study.

### Repetitive mild traumatic brain injury model

Beginning at ∼56PN, mice were subjected to either repeated sham or repeated closed head injury (CHI) delivered by the Impact One, Controlled Cortical Impact (CCI) Device (Leica; Wetzler, Germany) utilizing parameters which were previously described^3^. Mice were anesthetized with isoflurane (3.5% induction/2% maintenance) and fixed into a stereotaxic frame. Specifically, repeated CHI-CCI consists of 5 discrete concussive impacts to the head delivered at 24hr intervals generated by an electromagnetically driven piston (4.0m/s velocity, 3mm impact tip diameter, a beveled flat tip, 1.0 mm depth; 200 ms dwell time) targeted to bregma as visualized through the skin overlying the skull following depilation. Sham surgery consisted of identical procedures without delivery of impact. Body temperature was maintained at 37°C throughout by a warming pad and isoflurane exposure and surgery duration was limited to no more than 5 minutes. Following sham or CHI-CCI surgery completion, mice were immediately placed in a supine position in a clean cage on a warming pad and the latency to self-right was recorded.

### Recombinant adeno-associated viral vector injection

Seven-week-old mice were anesthetized with isoflurane (3.5% induction and 2% maintenance) and fixed into a stereotaxic frame. Body temperature was maintained at 37°C throughout the procedure and during recovery with a heating pad. The scalp was shaved and depilated with Nair ©, and a U-shaped incision between the ears was made on the skin overlying the skull, allowing for the visualization of bregma and lambda sutures during CHI-CCI procedure. Viral vectors were infused (50 nL/side; over 5min using a Nanoject III Injector, Drummond) using pulled glass pipettes into the LHb (coordinates from bregma: AP, −1.6 mm; ML, ± 0.5 mm; DV, −3.2 mm). Mice received either AAV8-CaMKII-hM4Di (Gi)-mCherry (Addgene# #50477) or control AAV8-CaMKII-EGFP (Addgene#50469) viral vector for viral expression in glutamatergic LHb neurons. Animals recovered for 10-11 days prior to the initiation of mTBI procedures. Viral expression was confirmed by fluorescence and/or immunohistochemistry at the conclusion of behavioral experiments and mice with no viral expression in the LHb were excluded from data analysis.

### Sucrose splash test

Sucrose splash test was performed at 10-12 days following the final mTBI or sham procedure in separate cohorts of sham and mTBI with or without viral injections. For chemogenetic inhibition of LHb activity, we utilized a novel DREADD ligand, JHU37160 ^28^ which has been shown to have high in vivo potency and no off-target effects. Mice that were injected with control or DREADD viral vectors received either 0.3mg/kg JHU37160 or equivalent volume of vehicle (0.9% saline) i.p. 30-min prior to behavioral testing. The order of drug presentation was cross-balanced within groups across behavioral testing days such that half of a group received vehicle and the other half received JHU37160. This dosage and the timing of administration/behavioral testing was chosen based on pharmacokinetic profile of JHU37160 indicating 30-min post-injection as the optimal time point where maximal drug concentration in brain is anticipated^28^. Mice were video monitored throughout the sucrose splash test. Mice were individually introduced to an empty (7×11.5×4.5 inches) clear polycarbonate cage. Following a 10-minute baseline video recording of behavioral activity the animal was removed from the testing arena, sprayed twice with an atomizer containing 10% sucrose solution onto the dorsal coat, returned to the test arena, and monitored for an additional 5 minutes. The 10% sucrose solution is a sticky substance that soils the animal’s coat, with the typical response being rapid initiation of vigorous grooming behaviors. Video recordings were assessed by an experimenter blinded to the condition of the subjects and scored for total grooming behavior and the latency to initiate the first bout of grooming after sucrose splash. Grooming is considered any movements involving active touching, wiping, scrubbing, or licking of the face, forelimbs, flank, or tail for greater than 3 consecutive seconds.

### Sucrose preference test

A separate cohort of sham and mTBI mice were tested for sucrose preference test 18 days following sham and mTBI surgeries. Mice were first single housed 7 days post-injury and left undisturbed for 12 days.

Starting on 12 days post-injury, all mice were presented with ad libitum access to two bottles containing either 1% sucrose solution or water to habituate the animals to the consumption of sucrose and avoid neophobia. After 3 days of habituation, mice returned to ad libitum access to drinking water only. On 18 days post-injury, mice were presented with two bottles at 1800 (start of their dark period); one bottle containing water and the other 1% sucrose solution, and evaluated for sucrose preference. Total consumption of each fluid was measured by weight the following day at 0600 (end of the dark period) and sucrose preference was calculated as the ratio of sucrose/total fluid consumption during the 12h test.

Throughout habituation days, bottles were counterbalanced daily to the left/right position to avoid a side-bias.

### Slice preparation

Mice were deeply anesthetized with isoflurane and immediately transcardially perfused with ice-cold artificial cerebrospinal fluid (aCSF) containing (in mM): 126 NaCl, 21.4 NaHCO_3_, 2.5 KCl, 1.2 NaH_2_PO_4_, 2.4 CaCl_2_, 1.00 MgSO_4_, 11.1 glucose, 0.4 ascorbic acid; saturated with 95% O_2_-5% CO_2_. Brain tissue was kept on ice-cold aCSF and tissue sections containing LHb were sectioned at 220μm using a vibratome (Leica; Wetzler, Germany) and subsequently incubated in aCSF at 34 °C for at least 1-hr prior to electrophysiological experiments. For patch clamp recordings, slices were then transferred to a recording chamber and perfused with ascorbic-acid free aCSF at 28-30 °C.

### Electrophysiology

Voltage-clamp cell-attached and voltage/current-clamp whole-cell recordings were performed from LHb neurons in sagittal slices containing LHb using patch pipettes (3-6 MOhms) and a patch amplifier (MultiClamp 700B) under infrared-differential interference contrast microscopy. Data acquisition and analysis were carried out using DigiData 1440A, pCLAMP 10 (Molecular Devices), Clampfit, Origin 2016 (OriginLab), and Mini Analysis 6.0.3 (Synaptosoft, Inc.). Signals were filtered at 3 kHz and digitized at 10 kHz.

To assess LHb spontaneous activity, cells were patch clamped with potassium gluconate-based internal solution (130 mM K-gluconate, 15 mM KCl, 4 mM adenosine triphosphate (ATP)-Na^+^, 0.3 mM guanosine triphosphate (GTP)-Na^+^, 1 mM EGTA, and 5 mM HEPES, pH 7.28, 275-280 mOsm) in slices perfused with aCSF. Spontaneous neuronal activity and AP firing patterns (tonic, bursting) were assessed in both cell-attached recordings in voltage-clamp mode at V=0 and whole-cell recording in current-clamp mode at I=0 for ∼1 min recording as previously described ^29, 30^. Spontaneous excitatory and inhibitory postsynaptic currents (sEPSCs and sIPSCs) were recorded within the same LHb neuron in voltage clamp mode with a cesium-methanesulfonate (CsMeS) based internal solution in intact synaptic transmission over 10 sweeps, each lasting 50 s (a total of 500s continuous recording for either sEPSC or sIPSC). Patch pipettes were filled with Cs-MeS internal solution (140 mM CsMeS, 5 mM KCl, 2 mM MgCl_2_, 2 mM ATP-Na+, 0.2 mM GTP-Na+, 0.2 mM EGTA, and 10 mM HEPES, pH 7.28, osmolality 275-280 mOsm) as previously described^31^. Cells were voltage-clamped at −55 mV to record spontaneous excitatory postsynaptic currents (sEPSCs) and +10 mV to record spontaneous inhibitory postsynaptic currents (sIPSCs) within the same neuron, as previously described^32^. The mean excitation/inhibition (sE/I) of spontaneous synaptic activity were calculated as sEPSC/sIPSC amplitude or frequency ratios from the same recording. The mean synaptic drive ratio was calculated as (sEPSC frequency × amplitude)/(sIPSC frequency × amplitude). To create histograms and cumulative probability plots for sE/I amplitude and frequency ratios and synaptic drive ratio, we adopted a novel quantitative approach ^33^ by randomly selecting 10 sEPSC and 10 sIPSC recordings (each lasting 15s) over each sweep of the 50s recording in each cell and calculated the ratios between sEPSC and sIPSC over 15s recordings. Therefore, for each cell, the combination of 10 sEPSC and 10 sIPSC yielded 100 data points (100 sE/I amplitude or frequency or synaptic drive ratios as calculated for the mean ratio values).

Whole-cell recordings of AMPAR-mediated miniature excitatory postsynaptic currents (mEPSCs) were isolated in aCSF perfused with picrotoxin (100µM), D-APV (50µM) and TTX (1 μM) and internal solution containing 117 mM Cs-gluconate, 2.8 mM NaCl, 5 mM MgCl_2_, 2 mM ATP-Na^+^, 0.3 mM GTP-Na^+^, 0.6 mM EGTA, and 20 mM HEPES (pH 7.28, 275–280 mOsm). Whole-cell recordings of GABA_A_R-mediated miniature inhibitory postsynaptic currents (mIPSCs) were isolated in aCSF perfused with DNQX; 10 μM, strychnine (1 μM) and tetrodotoxin (TTX, 1 μM). Patch pipettes were filled with 125 mM KCl, 2.8 mM NaCl, 2 mM MgCl_2_, 2 mM ATP-Na^+^, 0.3 mM GTP-Na^+^, 0.6 mM EGTA, and 10mM HEPES (pH 7.28, 275–280 mOsm). For both mIPSCs and mEPSCs, LHb neurons were voltage-clamped at −70 mV and recorded over 10 sweeps, each lasting 50 seconds. The cell input resistance and series resistance were monitored through all the experiments and if these values changed by more than 10%, data were not included.

### Statistics

Values are presented as mean ± SEM. The threshold for significance was set at *p < 0.05 for all analyses. All statistical analyses of data were performed using GraphPad Prism 8.4.1. For detecting the difference in distribution of silent, tonic or bursting LHb neurons in sham and mTBI mice, we used Chi-square tests. For the effects of mTBI on mean values of sEPSC, sIPSC, mEPSC, mIPSC, sE/I ratios, synaptic drive ratios, and sucrose splash test without viral injections, we used unpaired Student’s t test. Sucrose preference test results were analyzed by a non-parametric Mann-Whitney test. Mini Analysis software was used to detect and measure sEPSC and sIPSC amplitude and frequency (inter-event interval) as well as mEPSC and mIPSC amplitude, charge transfer (area under the curve), decay time constants (Tau) and frequency (inter-event interval) using preset detection parameters of spontaneous or mini events with an amplitude cutoff of 5 pA. Effects of mTBI on the cumulative probabilities data sets were analyzed using Kolmogorov-Smirnov tests. Effects of mTBI and JHU37160 during splash test were analyzed using 2- way ANOVA tests for with Sidak’s post-hoc test.

## Results

### mTBI increased spontaneous LHb tonic activity and excitatory synaptic drive onto LHb neurons

Here, we first investigated the effects of mTBI on spontaneous LHb activity in intact synaptic transmission in LHb slices from sham and mTBI male adult mice 2 weeks post-injury. We observed that mTBI increased the overall LHb spontaneous neuronal activity in both cell-attached voltage-clamp and whole-cell current-clamp recordings with intact synaptic transmission (Figure 1A-B, Chi squared test, p<0.05, p<0.01). Specifically, we found that in both types of recordings the percentage of neurons which were spontaneously active in tonic firing was larger in mTBI mice compared to control sham mice although we observed the opposite with bursting activity where fewer LHb neurons fired in bursting mode in mTBI mice compared to sham mice. Given that mTBI-induced increases in spontaneous excitatory synaptic drive onto LHb neurons could promote the overall LHb hyperactivity following mTBI, we next examined the effects of mTBI on spontaneous synaptic transmission in LHb neurons (Figures 1C-I, Figure 2). We measured both spontaneous excitatory and inhibitory postsynaptic currents (sEPSCs and sIPSCs) within the same LHb neurons while holding in the voltage clamp mode at −55 mV and +10 mV, respectively. We found that although the average amplitude and frequency of sEPSCs and sIPSCs were not significantly different between sham and mTBI mice, the cumulative probability amplitude and frequency (inter-event interval) plots of sEPSCs and sIPSCs were both significantly shifted. Specifically, there was an overall decrease in spontaneous excitatory and inhibitory synaptic transmission onto LHb neurons in mTBI mice compared to those from sham mice (Figure 2 A-D, Kolmogorov–Smirnov tests, p<0.0001). Spontaneous synaptic activity more likely reflects presynaptic effects of mTBI on glutamate and GABA release onto LHb neurons. While mTBI seems to decrease the overall synaptic transmission onto LHb neurons, it was possible that mTBI alters synaptic integration and shifts sE/I balance in LHb neurons. To determine sE/I ratios, we adopted a recent novel quantitative approach used for E/I calculations in parietal cortex layer 2/3 pyramidal cells given that a single measurement of mean E/I ratio in each cell cannot accurately reflect the dynamic balance between excitatory and inhibitory synaptic transmission within neurons that is constantly changing^33^. Based on this method, we generated 100 data points for both E/I amplitude and E/I frequency by including 10 segments of either sEPSC or sIPSC recordings (15sec) randomly selected over the period of 8-minutes of continuous recordings of either sEPSCs or sIPSCs and then plotted the histograms and cumulative probability plots of sE/I amplitude, sE/I frequency and synaptic drive ratios (Figure 1C-I). We found that although the mean sE/I amplitude and frequency in LHb neurons were not significantly different in mTBI and sham mice, mTBI resulted in significant shifts in cumulative probability curves of sE/I ratios (Figure 1 C-G, Kolmogorov–Smirnov tests, p<0.0001). Using the same data points for sEPSC and sIPSC amplitude and frequency, we also calculated and generated the mean, histogram, and cumulative probability curves of synaptic drive ratios in LHb neurons from sham and mTBI mice which incorporated all the properties of sEPSC and sIPSCs including amplitude, charge transfer and frequency^34^. While the mean synaptic drive ratios did not differ between sham and mTBI mice, mTBI resulted in a significant rightward shift in the cumulative probability curves of synaptic drive ratio suggesting that mTBI increased the overall excitatory synaptic drive onto LHb neurons (Figure 1H-I, Kolmogorov–Smirnov tests, p<0.0001). The majority of LHb neurons is believed to be glutamatergic and long-range projecting, although local glutamatergic and GABAergic connections within the LHb are reported ^18, 35-37^. Therefore, action potential-driven neurotransmitter release from LHb glutamatergic collaterals and intrinsic GABAergic interneurons also contributes to presynaptic effects of mTBI on glutamate and GABA release in sEPSC and sIPSC recordings. To independently examine the effects of mTBI on spontaneous, action potential-independent neurotransmitter release as well as postsynaptic function of AMPARs and GABA_A_Rs, we then recorded either mEPSCs or mIPSCs from LHb neurons from sham and mTBI mice at two weeks post-injury. mTBI significantly decreased the average charge transfer and Tau decay of mEPCSs and significantly shifted the cumulative probability curves of mEPSC frequency and tau decay (Figure 3, Student’s t-tests, Kolmogorov-Smirnov tests, p<0.05, p<0.0001). These data suggest that mTBI decreased the probability of presynaptic glutamate release onto LHb neurons from glutamatergic inputs and changed the kinetics of AMPAR mEPSCs. The cumulative probability curves of mIPSC amplitude and charge transfer were significantly shifted to the left following mTBI suggesting mTBI-induced decreases in postsynaptic GABA_A_R function, but interestingly mTBI also resulted in a significant leftward shift in the cumulative probability curve of frequency (inter-event interval) that suggests possible increases in the probability of presynaptic GABA release from a subset of GABAergic inputs onto LHb neurons (Figure 4, Kolmogorov-Smirnov tests, p<0.0001).

**Figure 1.**
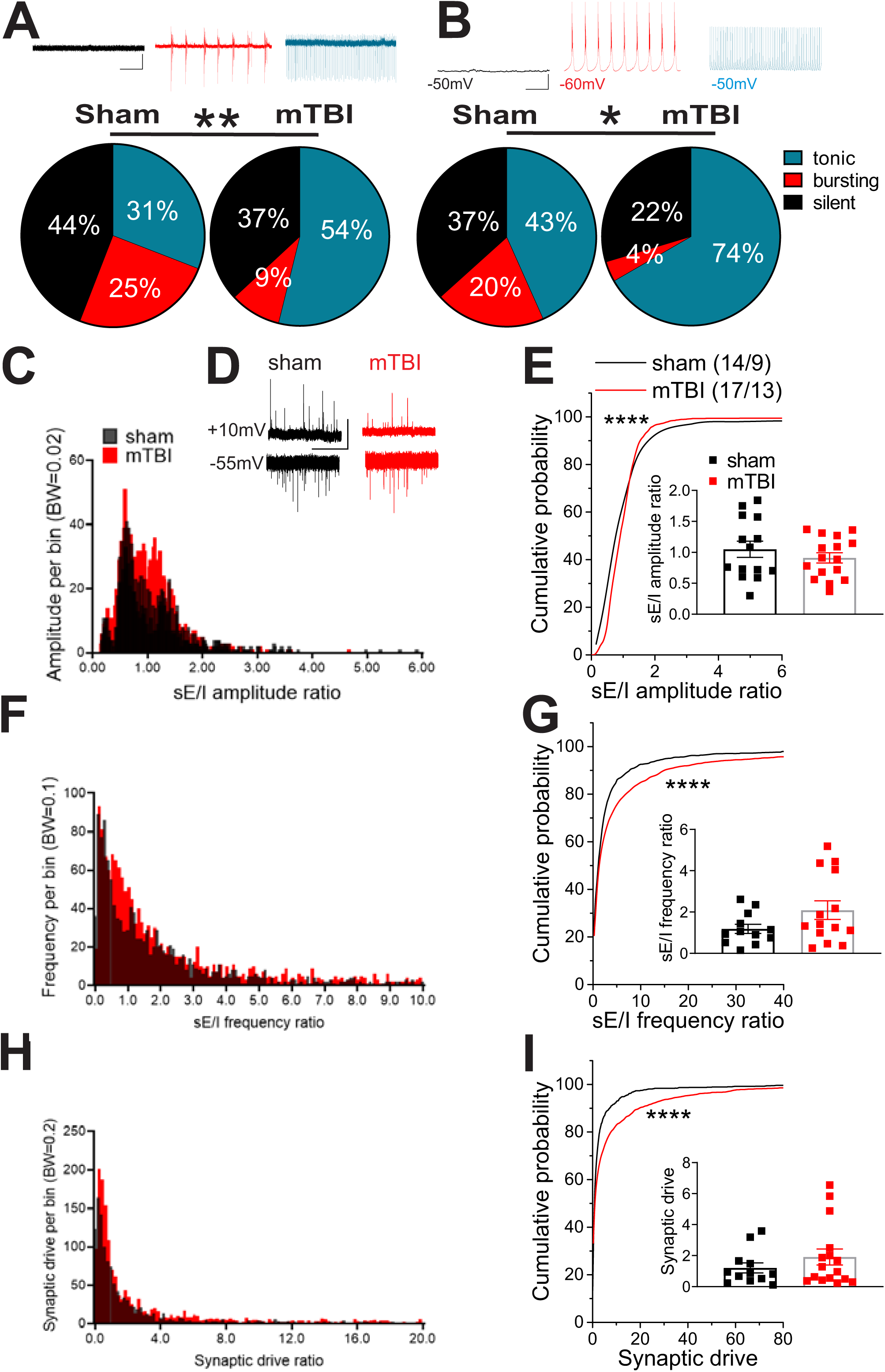
Effects of mTBI on LHb spontaneous neuronal and synaptic activity. Pie charts and representative traces of voltage-clamp cell-attached recordings (**A**, V=0 mV, sham, n=68/11; mTBI, n=65/14) and current-clamp whole-cell recordings (**B**, I=0 pA, sham, n=30/11; mTBI, n=27/14) of spontaneous neuronal activity across sham and mTBI mice. Comparison of the percent distributions of silent (black), tonic (blue), or bursting (red) LHb neurons increased their tonic LHb neuronal activity while decreased their bursting activity following mTBI. **C, F** and **H** show the histograms of spontaneous excitatory/inhibitory (sE/I) amplitude, frequency and synaptic drive ratios in LHb neurons recorded from sham and mTBI mice. **D** shows representative voltage-clamp recordings of spontaneous excitatory postsynaptic currents (sEPSCs, recorded at −55mV) and spontaneous inhibitory postsynaptic currents (sIPSCs, recorded at +10 mV) within the same LHb neurons in sham (top, black) and mTBI (bottom, red) mice (calibration bars, 50pA/5 s). **E, G** and **I** show the average and cumulative probability plots of the sE/I amplitude, frequency and synaptic drive in LHb neurons recorded from sham and mTBI mice. mTBI significantly resulted in significant shifts in the distribution curves of sE/I amplitude, frequency and synaptic drive ratios, resulting in an overall increased excitatory synaptic drive in LHb neurons. *p<0.05, **p<0.01, ****p<0.0001 by Chi squared or Kolmogorov–Smirnov tests. n represents the number of recorded cells/mice.

**Figure 2.**
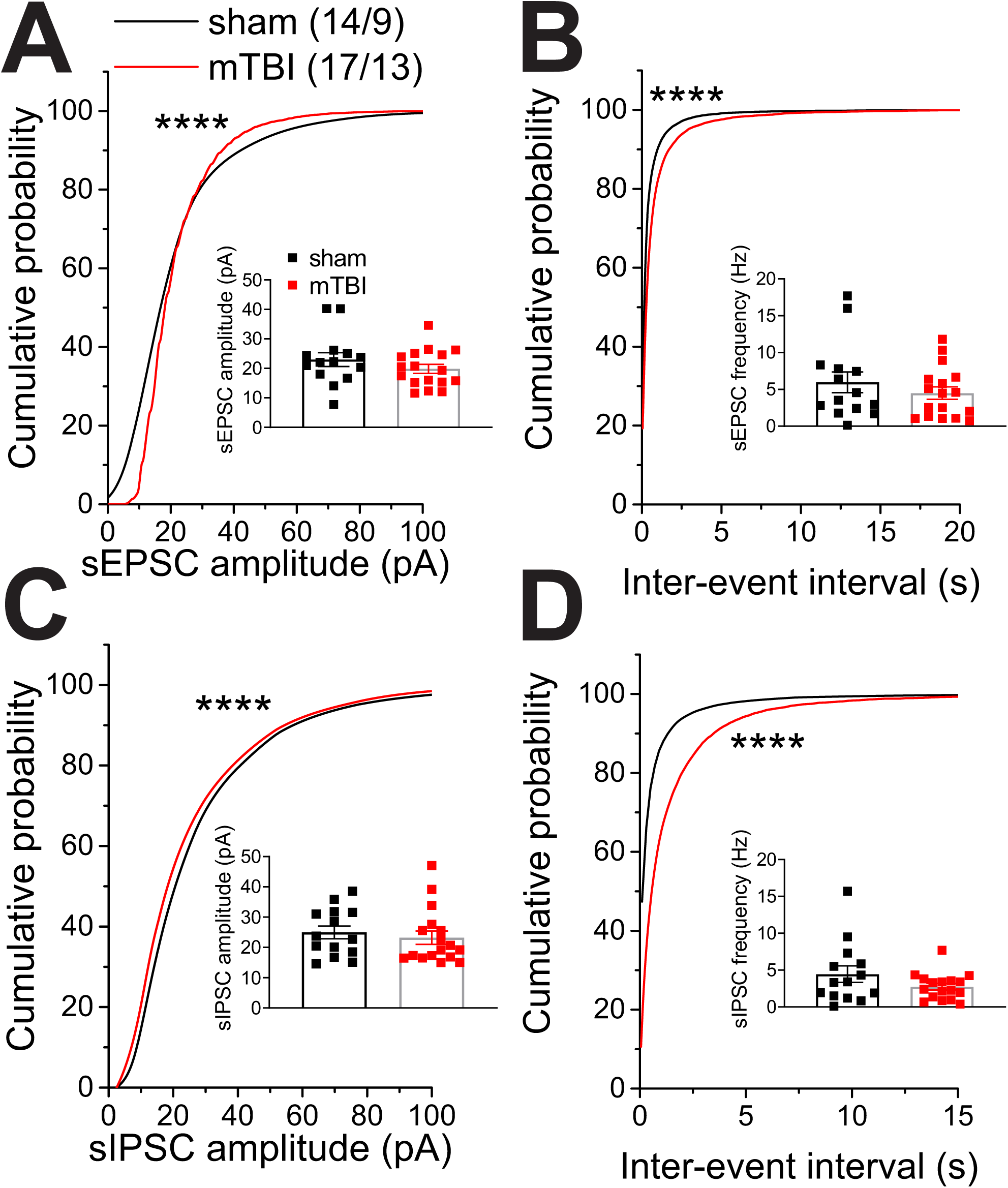
Effects of mTBI on spontaneous synaptic activity in LHb neurons. Graph shows average and cumulative probability amplitude and frequency (inter-event interval) plots of spontaneous excitatory postsynaptic currents (sEPSCs, recorded at −55mV, **A-B**) and spontaneous inhibitory postsynaptic currents (sIPSCs, recorded at +10 mV, **C-D**) within the same LHb neurons from sham and mTBI mice at two weeks following the injury. mTBI significantly shifted cumulative probability curves of sEPSC and sIPSC amplitude and frequency, resulting in an overall decreased spontaneous excitatory and inhibitory transmission in LHb neurons. ****p<0.0001 by Kolmogorov–Smirnov tests. “n” in this and all following graphs represents the number of recorded cells/mice.

**Figure 3.**
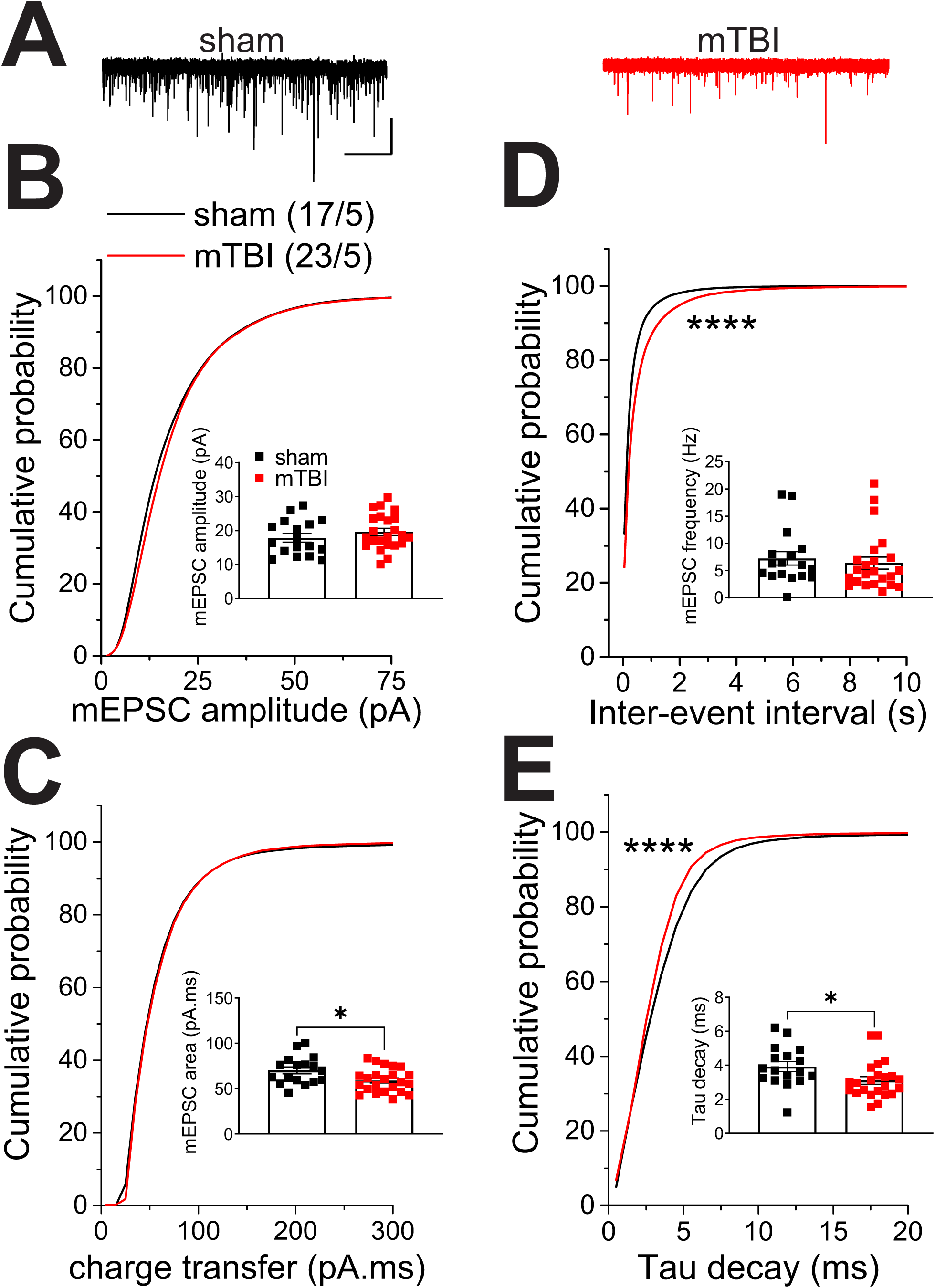
Effects of mTBI at LHb glutamatergic synapses. **A**) Representative AMPAR-mediated miniature excitatory postsynaptic current (mEPSC) traces from sham (black) and mTBI (red) mice (calibration bars, 30pA/5 s), average and cumulative probability plots of mEPSC **B**) amplitude, **C**) charge transfer (area under the curve), **D**) frequency (inter-event interval) and **E**) decay time constants (Tau) in sham and mTBI mice at two weeks following the injury. mTBI significantly decreased the average charge transfer and Tau decay of mEPCSs and shifted the cumulative probability curves of mEPSC frequency and tau decay, suggesting mTBI-induced decreases in the probability of presynaptic glutamate release onto LHb neurons and changes in the kinetics of AMPAR mEPSCs. *p<0.05, ****p<0.0001 by unpaired Student’s t-tests or Kolmogorov–Smirnov tests.

**Figure 4.**
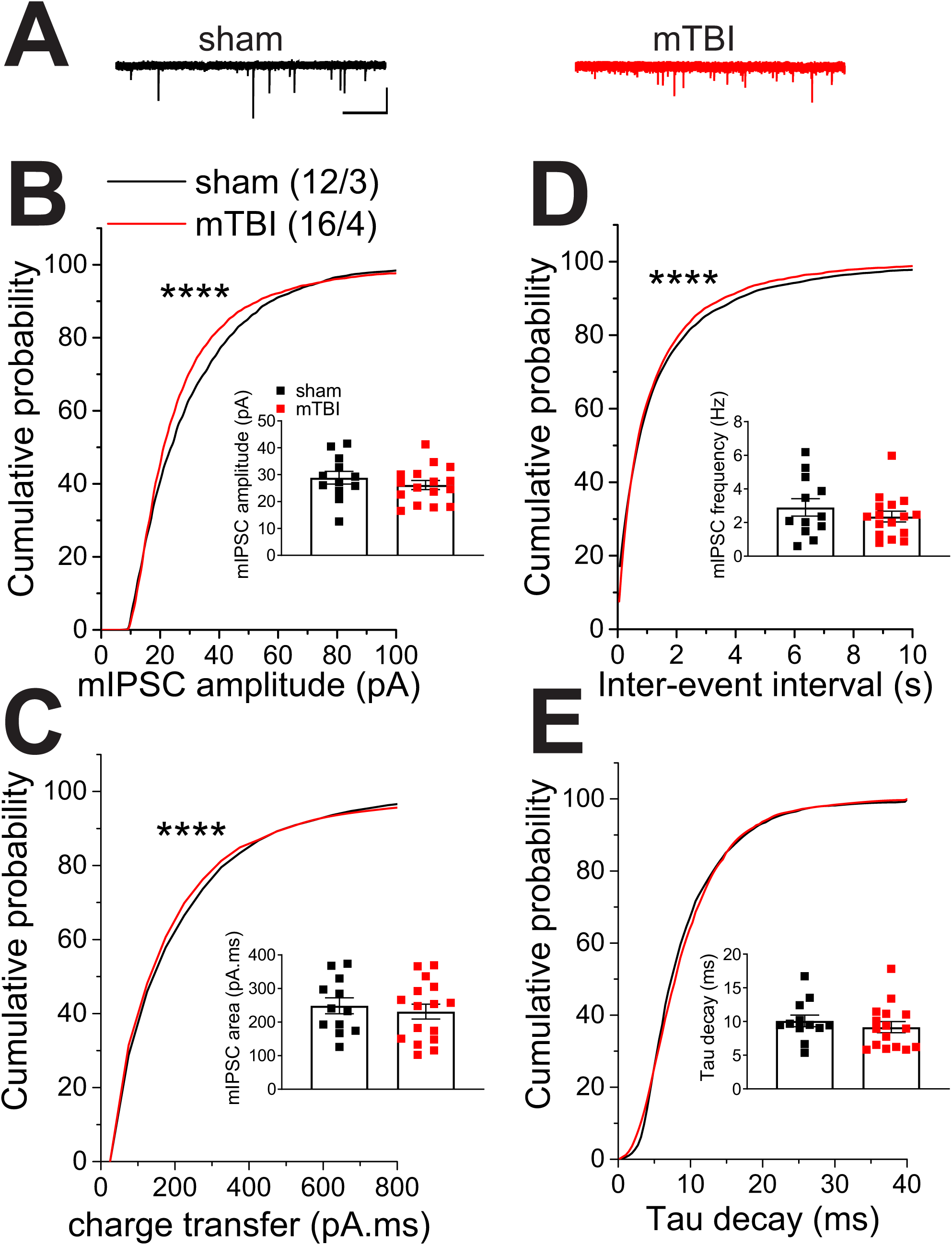
Effects of mTBI at LHb GABAergic synapses. **A**) Representative GABA_A_R-mediated miniature inhibitory postsynaptic current (mIPSC) traces from sham (black) and mTBI (red) mice (calibration bars, 50pA/5 s), average and cumulative probability plots of mIPSC **B**) amplitude, **C**) charge transfer (area under the curve), **D**) frequency (inter-event interval) and **E**) decay time constants (Tau) in sham and mTBI mice at two weeks following the injury. mTBI significantly shifted the cumulative probability curves of mIPSC amplitude, charge transfer and frequency, suggesting mTBI-induced decreases in postsynaptic GABA_A_R function accompanied by increases in the probability of presynaptic GABA release onto LHb neurons. ****p<0.0001 by Kolmogorov–Smirnov tests.

### Chemogenetic inhibition of LHb glutamatergic neurons reversed mTBI-induced deficits in sucrose splash test

Previously, it has been shown that this model of mTBI is associated with social deficits including reduced social interactions at three weeks following the injury ^3^. To test whether mTBI also induces other negative affective states in addition to social anhedonia that could be indicative of depressive phenotype, we conducted sucrose splash test (an index for motivated self-care behavior) and sucrose preference test (an index for sucrose anhedonia) in sham and mTBI mice at 10-18 days post-injury (Figure 5 A-C). We found that mTBI significantly increased the latency to self-grooming without an overall change in total grooming time in sucrose splash test but did not affect sucrose preference in sucrose preference test (Figure 5 A-C, Student’s t-tests, t=3.31, df=15, p<0.01). To provide a causal link between LHb hyperactivity and motivational deficits in sucrose splash test, we then employed a chemogenetic approach using a novel DREADD agonist, JHU37160. Two weeks prior to sham or mTBI procedures, mice were injected bilaterally into the LHb with either control virus (AAV-CamKII-eGFP) or inhibitory DREADD (AAV-CamkIIa-hM4D (Gi)-mCherry, Gi-DREADD). We verified the effectiveness of chemogenetic manipulation of LHb activity in electrophysiological recordings with bath application of JHU37160 which maximally inhibited LHb neuronal excitability at 30min post-application in Gi-DREADD- expressing LHb neurons (Figure 6A-B) but not control virus (data not shown). Behavioral testing was then performed two weeks following sham and mTBI procedure (four weeks after the viral injections allowing for optimal expression for DREADD constructs). In control virus-injected groups, the main effect of mTBI on increasing grooming latencies in sucrose splash test remained significant with no change in total time spent for self-grooming, while JHU37160 also did not affect self-grooming behaviors in both sham and mTBI mice injected with control virus (Figure 5D-E, two-way ANOVA tests; effect of mTBI: F (1, 15) = 8.43, p<0.05). In Gi-DREADD-injected groups, mTBI significantly increased grooming latencies with no change in total grooming time with saline injections. Chemogenetic inhibition of LHb glutamatergic neurons with JHU37160 normalized self-grooming behaviors in mTBI mice with a return of grooming latencies comparable to those of sham mice (Figure 6C-D, two-way ANOVA tests; effect of mTBI: F (1, 27) = 13.21, effect of JHU37160: F (1, 27) = 5.15, mTBIxJHU37160 interaction: F (1, 27) = 4.764, p<0.01, p<0.05).

**Figure 5.**
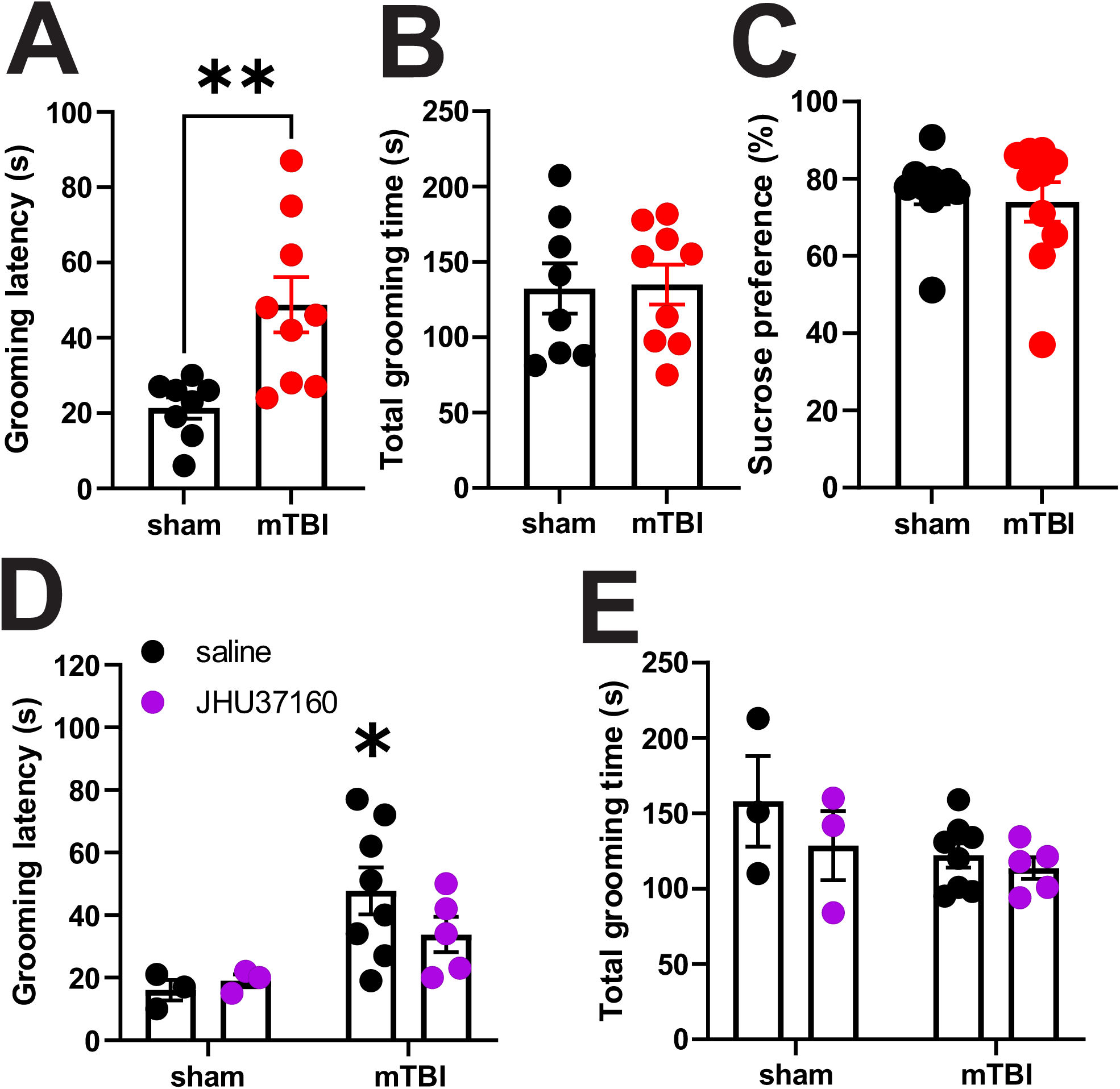
Effects of mTBI on sucrose splash and preference tests. **A-B**) mTBI significantly increased the latency to grooming without an overall change in total grooming time in sucrose splash test. **C**) mTBI did not alter sucrose preference in sucrose preference test. **D-E**) Sham (n=3/group) and mTBI (n=7- 8/group) mice injected with control virus (AAV-CamKII-eGFP) into the LHb 4 weeks prior behavioral testing, received i.p. injections of saline or 0.3mg/kg JHU37160 30min before sucrose splash tests. Latencies to grooming and total grooming time were measured following sucrose splash in sham and mTBI control mice. mTBI increased grooming latencies with no effects of JHU37160 on behavior in sham or mTBI control mice.

**Figure 6.**
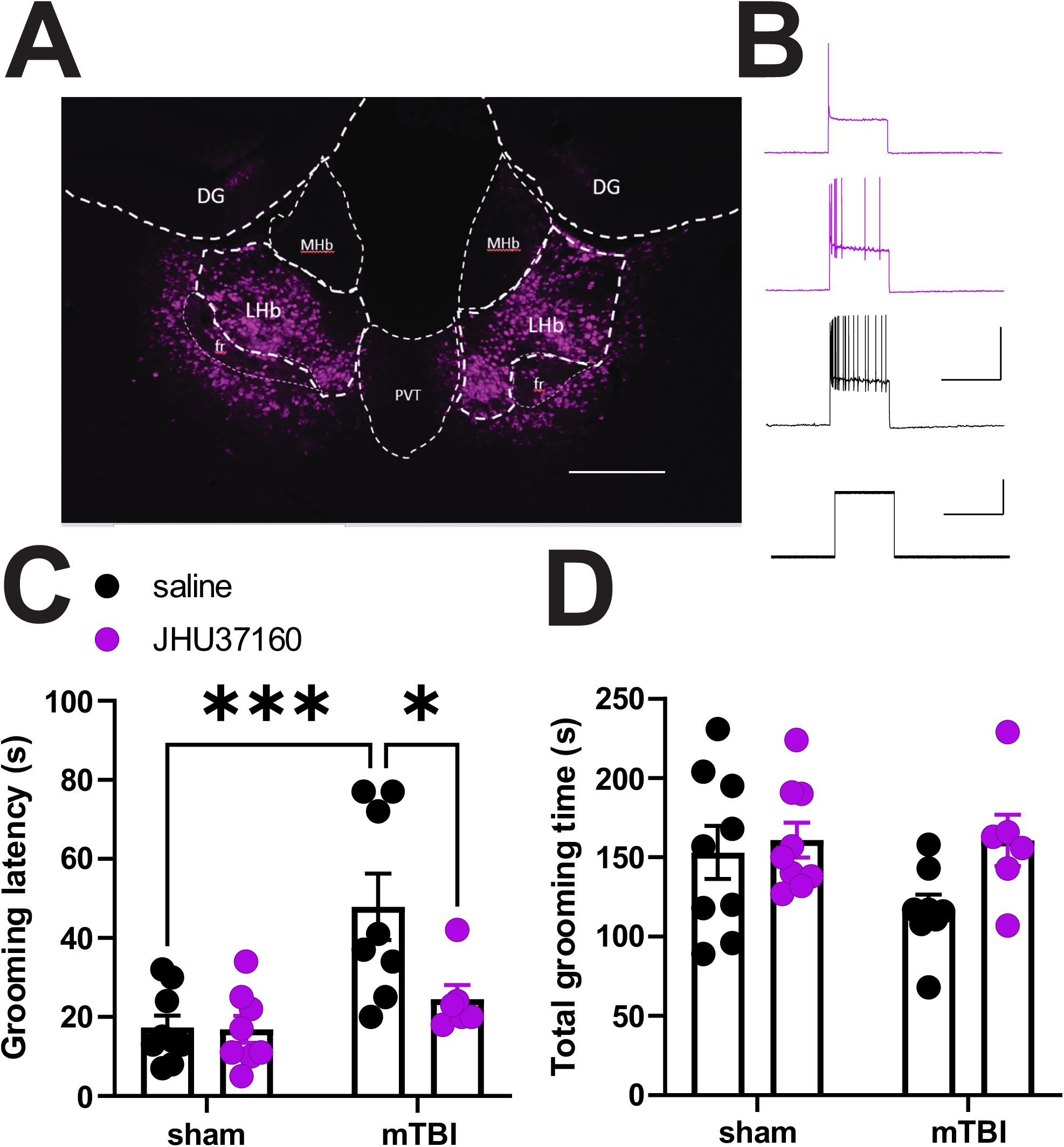
Effects of chemogenetic inhibition of LHb glutamatergic neurons in sucrose splash test in sham and mTBI mice. **A** shows representative image of a coronal section of LHb with bilateral injection of AAV-CamkIIa-hM4D (Gi)-mCherry (Gi-DREADD, purple) in the LHb. **B** shows sample current clamp recording of action potentials from a Gi-DREADD expressing LHb glutamatergic neuron at baseline, 15min and 30 min after bath application of the DREADD-specific agonist, JHU37160 (100nM), in LHb slice from a Gi-DREADD mouse demonstrating the effectiveness of the Gi-DREADD strategy in chemogenetic inhibition of LHb glutamatergic neurons. **C-D** show sham (n=8-9/group) and mTBI (n=6- 8/group) mice injected with Gi-DREADD into the LHb that received i.p. injections of either saline or 0.3mg/kg JHU37160 30min before sucrose splash tests. Latencies to grooming and total grooming time were measured following sucrose splash in sham and mTBI Gi-DREADD mice. mTBI-induced increases in grooming latencies were reversed by chemogenetic inhibition of LHb glutamatergic neurons. Two-way ANOVA, * p<0.05, **p<0.01.

## Discussion

Here, we used a mouse model of mTBI which incorporates repetitive closed head injuries that is relevant to the most common form of mTBI in humans (i.e., repeated and mild concussive head injuries)^3^. We provided evidence that mTBI-induced alterations in synaptic integration promotes LHb tonic activity which contributes to motivational deficits in self-care and grooming behavior in adult male mTBI mice. Contrary to many preclinical studies using an open skull injury model of mTBI, our model utilized a closed head injury avoiding opening of the skull, which itself may provoke considerable damage and changes in immune function in the brain ^38^. Although our model and other mTBI animal models may not predict all outcomes related to mTBI in humans, we observed that this mTBI model is associated with motivational deficits in self-care grooming behavior in addition to the previously reported social anhedonia associated with this model ^3^ in male mice suggesting the possibility of mTBI-induced reward/motivational circuit dysfunction underlying long-term negative affective states of mTBI. The cortical region under the impact site used in our model includes the mPFC, an important integrative hub involved in processing of a wide range of cognitive, sensory, social, motivational and emotional-related information. Specifically the anterior cingulate cortex (ACC, which is considered the dorsal component of mPFC ^39^) exhibit low levels of axonal damage in this mTBI model^3^. Therefore, it was likely that mTBI alters the function of subcortical regions which receive direct^40-45^ and indirect^39, 46-50^ projections from mPFC and ACC, such as the LHb, a critical stress- and reward-related brain region that promotes anhedonia and motivational deficits. We observed that tonic firing LHb neurons were more prevalent following mTBI which is consistent with the literature that LHb hyperactivity in general contributes to the development of depression-like motivational and social deficits, and anhedonic phenotypes^2, 20, 24, 29, 51-53^. Intriguingly, mTBI was associated with fewer LHb bursting neurons. This is in contrast with the most common finding of preclinical models of depression where NMDA receptor-dependent LHb bursting are also elevated^19, 24, 29, 30^ and is shown to be the critical target for anti-depressant effects of ketamine^19, 24^. mTBI may result in alteration of synaptic integration due to an imbalance of E/I that could explain changes in LHb activity and firing patterns. Consistent with this idea, we found that while mTBI significantly reduced the overall spontaneous excitatory glutamatergic and GABAergic synaptic transmission onto LHb neurons, the greater suppression of GABAergic transmission resulted in a significant shift of E/I and synaptic drive ratios toward excitation which supports the enhanced tonic activity of LHb. Additionally, diminished GABAergic inhibition could also mediate deficits in LHb bursting given that hyperpolarization is needed to trigger LHb bursting activity ^19^. mTBI similarly affected sEPSC and mEPSC recordings suggesting a global reduction in presynaptic glutamate release from glutamatergic inputs to the LHb, but this was not the case for GABAergic inputs. Both sIPSC and mIPSC measurements of amplitude and charge transfer were indicative of a significant reduction in postsynaptic GABA_A_R function by mTBI. In contrast, mTBI exerted opposite shifts in sIPSC and mIPSC inter-event interval cumulative probability curves, such that sIPSC recordings suggest a significant decrease in presynaptic GABA release, whereas mIPSC recording indicated a slight increase in presynaptic GABA release. This seemingly confounding result may be due to the integration of glutamatergic excitatory collaterals within the LHb and intrinsic LHb GABAergic neurons^18, 35, 36, 54^. LHb GABAergic interneurons expressing parvalbumin (PV) inhibit local LHb neurons ^18^. Additionally, LHb GABAergic neurons expressing estrogen receptors regulate motivated behaviors, mostly projecting locally within the LHb, although a few long-range projections to midbrain (RMTg, substantial nigra pars reticulata) have been observed ^35^. Recently, it has been shown that a small subpopulation of LHb GABAergic neurons that express glutamic acid decarboxylase 2 (GAD2) and orexin receptor 2 (OxR2) can be activated by LH orexin neurons that project to the LHb. Their activation results in an overall inhibition of LHb activity and promotes male–male aggression and conditioned place preference for intruder-paired contexts in mice ^54^. Given that in mIPSC recordings action potential-driven neurotransmitter release from local inhibitory inputs are blocked by TTX, the difference shown between sIPSC and mIPSC recordings indicates that mTBI-induced suppression of local LHb GABAergic neurons may significantly dampen this intrinsic inhibitory drive within the LHb microcircuit. Collectively, altered intrinsic inhibitory drive could be responsible for mTBI-induced E/I shifts to excitation that promotes LHb tonic activity. On the other hand, mTBI seems to increase presynaptic GABA release from a subset of extrinsic GABAergic inputs to LHb that became unmasked in mIPSC recordings. It is possible that this slight increase is a homeostatic response to the overall mTBI-induced LHb hyperactivity. mTBI also induced a change in AMPAR decay kinetics (decreases in Tau) suggesting a possible accumulation of GluA2 lacking AMPARs [i.e., calcium-permeable (CP)-AMPARs] at glutamatergic synapses onto LHb neurons following mTBI. As mentioned earlier, mPFC and ACC cortical areas provide direct projections to LHb ^40-45^ as well as widespread direct and indirect projections to other cortical and subcortical brain regions including LH, VTA, EP, NAc, CeA, ventral pallidum (VP) ^39, 46-50, 55^; all of which provide a direct input to LHb neurons ^16, 45, 47, 56-60^. It has been shown that EP and LH glutamatergic transmission onto VTA-projecting LHb neurons are mainly mediated by CP-AMPARs that display strong inward rectification and have faster decay kinetics^53, 61^. Therefore it is possible that the increase in insertion of CP-AMPARs by mTBI becomes restricted to these specific synapses onto VTA-projecting LHb subpopulation. Our observation that chemogenetic inhibition of LHb glutamatergic neurons reversed motivational deficits supports the idea that the general LHb hyperactivity due to the altered synaptic integration by mTBI plays a causal role in reducing self-care motivated behavior in mTBI mice.

Chemogenetic inhibition of DRN-projecting LHb neurons is sufficient to decrease immobility in the forced swim test (as a measure of behavioral despair) and reduces behavioral flexibility of mice to adjust effort to obtain saccharin reward in saccharin two bottle choice licking behavior when saccharin was omitted while chemogenetic activation of DRN-projecting LHb neurons increases the effort of mice under this frustrating conditions ^62^. Overall, this study suggests that hyperactivity of DRN-projecting LHb neurons can mediate anhedonic states under heightened stress and decreases the behavioral flexibility in adaptive coping behaviors when the conditions change, therefore it is possible that strengthening of DRN- projecting LHb projections play a role in motivational deficits following mTBI. However, chemogenetic inhibition of DRN-projecting LHb neurons decreases social interaction in three-chambered social preference test in male mice ^63^ suggesting that this pathway may not mediate mTBI-induced social deficits. Interestingly, behavioral avoidance from social interaction can be triggered after stimulation of LHb activity by PFC inputs in rats ^44^. Moreover, optogenetic stimulation of mPFC-projecting LHb neurons during the forced swim test in rats decreases the frequency of kicks (as a measure of active escape behavior) which corresponds to increases in immobility (passive coping behavior)^43^. This may suggest that rather than direct mPFC projections to LHb, the indirect mPFC projections through other LHb-projecting areas are involved in behavioral deficits following mTBI. Given the complexity of LHb upstream and downstream circuits in mediating different aspects of motivated and affective behaviors, it would be worthwhile to employ intersectional DREADD-based approach combined with optogenetics and electrophysiology to investigate causal roles of distinct intrinsic and extrinsic inputs to LHb and related LHb subpopulations and synaptic adaptations in mTBI-induced social and motivational deficits.

## Conclusions

Our study suggests that mTBI can dysregulate reward and motivational circuit function at the level of subcortical structures such as the LHb that plays an important role in decision making and behavioral flexibility under stressful and aversive conditions. Our current mTBI model offers a valid repeated concussion model for investigation of potential therapeutic circuit interventions for prevention and treatment of psychiatric disorders in patients with a history of repetitive mTBI.

## Acknowledgments

The opinions and assertions contained herein are the private opinions of the authors and are not to be construed as official or reflecting the views of the Uniformed Services University of the Health Sciences or the Department of Defense or the Government of the United States. The authors would like to thank Dr. Veronica Alvarez at NIAAA for their invaluable support in establishing chemogenetic studies at Nugent Laboratory. Behavioral testing and analysis were performed in the Preclinical Behavior and Modeling Core at the Uniformed Services University.

## Authorship contribution

FN and WF were responsible for the study concept and design. WF, LL, MR, SS, SG and MT contributed to the acquisition of animal data. FN, WF, LL, MR, SS, SG, MT, BC and AS assisted with data analysis and interpretation of findings. FN, WF, LL, MR, SS and MT wrote the initial draft of the manuscript. All authors critically reviewed the content and approved final version of manuscript for submission. The authors acknowledge Dr. Yeonho Kim, Dr. Amanda Fu and Laura Tucker at the USUHS Preclinical Modeling and Behavior Core for supporting the studies.

## Data Sharing

The data that support the findings of this study are available on request from the corresponding author. The data are not publicly available due to privacy or ethical restrictions.

